# Rescue of stalled clathrin-mediated endocytosis by asymmetric Arp2/3-mediated actin assembly

**DOI:** 10.1101/2021.07.16.452693

**Authors:** Meiyan Jin, Cyna Shirazinejad, Bowen Wang, Amy Yan, Johannes Schöneberg, Srigokul Upadhyayula, Ke Xu, David G. Drubin

## Abstract

Actin assembly facilitates vesicle formation in several trafficking pathways. Clathrin-mediated endocytosis (CME) shows elevated actin assembly dependence under high membrane tension. Why actin assembly at CME sites occurs heterogeneously even within the same cell, and how assembly forces are harnessed, are not fully understood. Here, endocytic dynamics, actin presence, and geometry of CME proteins from three different functional modules, were analyzed using three-dimensional (3D) super-resolution microscopy, live-cell imaging, and machine-learning-based computation. When hundreds of CME events were compared, sites with actin assembly showed a distinct signature, a delay between completion of coat expansion and vesicle scission, indicating that actin assembly occurs preferentially at stalled CME sites. N-WASP is recruited to one side of CME sites where it is positioned to stimulate asymmetric actin assembly. We propose that asymmetric actin assembly rescues stalled CME sites by pulling vesicles into the cell much like a bottle opener pulls off a bottle cap.

## Introduction

Formation of clathrin-coated vesicles requires forces to first bend the membrane into a sphere or tube, and to then break the thin neck that connects the vesicle to the plasma membrane. These forces are generated through the combined actions of actin filament assembly and proteins that directly bend the membrane^1–5^ (Supplementary Fig. 1a). Several studies have demonstrated that dependence of CME on actin assembly increases under elevated membrane tension^6–9^. Interestingly, actin does not assemble at all CME sites in mammalian cells, suggesting highly localized differences in requirement for actin assembly, that nature of which are obscure^10–12^. A detailed understanding of how actin forces are harnessed to aid vesicle formation and scission, and whether and how actin assembly might mediate an adaptive response to the opposing forces such as membrane tension and turgor pressure, depends on understanding which CME sites assembly actin, where filament assembly occurs around the endocytic membrane and when. In yeast cells, where turgor pressure is particularly high, super-resolution data suggest that actin assembles symmetrically around CME sites and indicate that actin regulators including Las17, which is yeast WASP, are present in a ring surrounding the base of the clathrin coat symmetrically^13^. On the other hand, studies on fixed mammalian cells raised the possibility that actin assembly may at least in some cases be initiated asymmetrically at clathrin coats^14,15^. However, methods used for these studies prevented analysis of large numbers of sites, and suffered from possible loss of actin filaments during unroofing and extraction of the cells. Which CME sites assemble actin, and how actin networks are organized with respect to CME sites, has not been determined systematically, in a large-scale, unbiased manner, particularly in live mammalian cells. This information is essential to understanding how and why actin assembly forces are harnessed for CME.

Here, by combining fixed and live-cell imaging of triple-genome-edited, human induced pluripotent stem cells (iPSCs), and newly developed machine-learning-based computational analysis tools, we report that N-WASP and Arp2/3 complex localize at one side of the coat and neck of invaginating endocytic sites until the scission, similar to what was proposed previously^14,15^. Most importantly, by comparing recruitment dynamics of proteins from three distinct endocytic modules for over one thousand endocytic events, we found that branched actin assembly occurs predominantly at sites that have stalled between coat expansion and scission. We propose that these branched actin networks rescue stalled CME.

## Results

### Super-resolution imaging reveals asymmetric actin distribution around endocytic sites

To investigate the physiological roles and spatiotemporal regulation of actin assembly at CME sites in mammalian cells, we applied genome-editing techniques to generate a human iPSC line (hereafter referred to as ADA cells) that co-expresses a TagRFP-T fusion of the mu1 subunit of the AP2 adaptor complex (AP2M1), a TagGFP2 fusion of dynamin2 (DNM2), and a HaloTag fusion of the ARPC3 subunit of the Arp2/3 complex as representatives of the CME coat, scission and actin modules respectively^2,16,17^ (Supplementary Fig. 1 and Supplementary Video 1, 2). Previous studies showed that endogenously tagged AP2M1, DNM2 and ARPC3 can serve as reliable markers of these CME functional modules that avoid disruption of physiological spatiotemporal organization of the process as might be caused by overexpression of fluorescently labeled proteins^12,18–21^. We observed dynamic CME events on the basal plasma membrane of the genome-edited cells using Total Internal Reflection Fluorescence (TIRF) microscopy (Supplementary Fig. 1c). Consistent with previous studies, AP2 is recruited at early CME stages while DNM2 is recruited in two phases^10,16,20,22^. At the early stage of CME, a relatively small amount of DNM2 is recruited to CME sites. Shortly before the end of a CME event, the DNM2 recruitment rate increases rapidly with DNM2 levels reaching a peak concomitant with vesicle scission^16,20,23^ (Supplementary Fig. 1c). This later rapid-recruitment phase represents the assembly of the dynamin helix on the highly curved neck of the budding vesicle after the U to Ω shape transition of the endocytic membrane^20,23–27^.

Super-resolution imaging of fixed human skin melanoma SKMEL cells observed asymmetry of actin arrangement around CME sites^7^, consistent with observations from previous studies^14,15^. To analyze how actin networks are organized at CME sites in iPS cells, we first performed two-color 3D Stochastic Optical Reconstruction Microscopy (STORM) imaging^28^ on fixed ADA cells, localizing either AF647 phalloidin-labeled actin filaments^29^ or HaloTag-fused ARPC3 at CME sites. Due to the dense phalloidin labelling of cortical actin filaments under the plasma membrane, it was often challenging to unambiguously identify the CME-specific actin structures in iPSCs. However, in regions with thinner cortical actin layers, we observed that actin was typically distributed asymmetrically around CME sites (Fig. 1a, b), consistent with what has been observed in different mammalian cell lines by STORM or EM imaging approaches^7,15^. Antibody labeling of ARPC3-Halotag in the ADA cells had the advantage of a less complex staining pattern. Besides being highly concentrated in lamellipodia, ARPC3 was associated with CME sites asymmetrically, like actin (Fig 1c, d). These data suggest an asymmetric Arp2/3-mediated actin network arrangement around CME sites.

**Fig. 1:**
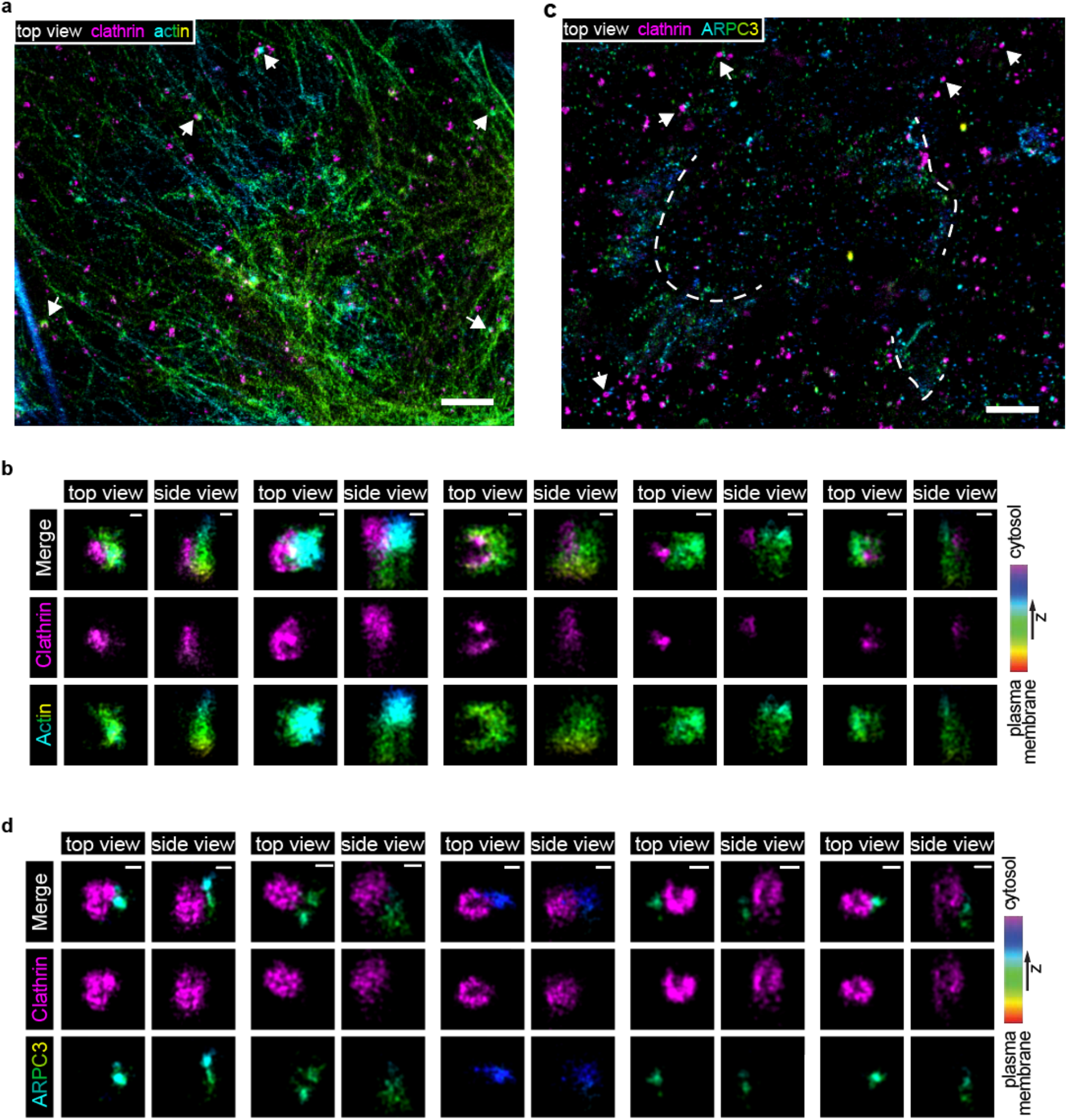
Two-color, 3D stochastic optical reconstruction microscopy (STORM) shows that actin structures are off-centered with respect to clathrin coats. **a, b**, Two-color 3D STORM image of bottom membrane of ADA cells immunolabeled with clathrin light chain antibody (clathrin, CF-680, magenta) and phalloidin (actin, AF647, rainbow). **c, d**, Two color 3D STORM image of the bottom membrane of ADA cells immunolabeled with clathrin light chain antibody (clathrin, AF647, magenta) and HaloTag antibody (ARPC3-HaloTag, CF-680, rainbow). Dotted lines label lamellipodia. **b, d**, The highlighted CME sites, which are labeled by white arrows in (**a**) and (**c**), are rotated and shown in magnified top and side view projections. Color bar shows the z position of ARPC3-HaloTag. Scale bars: 2µm, 100nm.

### Asymmetric branched actin networks assembled at CME sites persist through scission

We next used ADA cells to investigate actin assembly at CME sites in live cells, which has several advantages over studies in fixed cells. During the fixation and subsequent sample preparation, actin structures may not be faithfully preserved. In addition, in fixed cells it is very difficult to identify the stage of the CME, so the timing, geometry and dynamics of actin assembly cannot be related to the endocytic stage. More importantly, only by using live cells is it possible to trace a single CME event from start to finish, and to therefore identify those CME events wherein no detectable actin is ever assembled so key parameters can be compared between events with and without associated actin assembly.

By visualizing endogenously tagged AP2M1 to mark the coat and CME initiation, and DNM2 to mark the neck and scission, together with ARPC3 to specifically label Arp2/3-nucleated, branched actin filaments (Supplementary Fig. 1a), we were able to precisely study the spatial and temporal regulation of actin assembly during CME. Three-color labeling and analysis of the displacement between markers for the three modules allowed us to distinguish *bona fide* asymmetric actin assembly from events that artificially might appear asymmetric because the invaginations were elongated and tilted (Fig2 a). Using TIRF live-cell imaging, we observed ARPC3-labeled branched actin networks at lamellipodia and a subpopulation of CME sites (Fig. 2b, c). Dynamic actin assembly and disassembly occurred at CME sites with different spatio-temporal characteristics, including discrete CME sites, clathrin plaques and at clathrin coat splitting sites, as previously reported^14^ (Fig. 2d and Supplementary Fig. 2a, b). In the analysis described below, we focus on the discrete CME events and not the more complex ones (plaques and splitting events). Analysis of these events with 1s/frame temporal resolution revealed that ARPC3 is most robustly recruited during the late stages of CME shortly before scission^12^ (Fig. 2c, d). Interestingly, we observed clear spatial displacement between ARPC3 (actin module) and AP2 (coat module) as well as between ARPC3 and DNM2 (neck) before vesicle scission (Fig. 2d). This observation supports the conclusion that asymmetric branched actin networks provide forces at endocytic sites through the time of scission. Imaging fluorescent beads using the same settings indicates that the displacement is not an artifact caused by misalignment between different imaging channels (Supplementary Video 3 and Supplementary Fig. 2c).

**Fig. 2:**
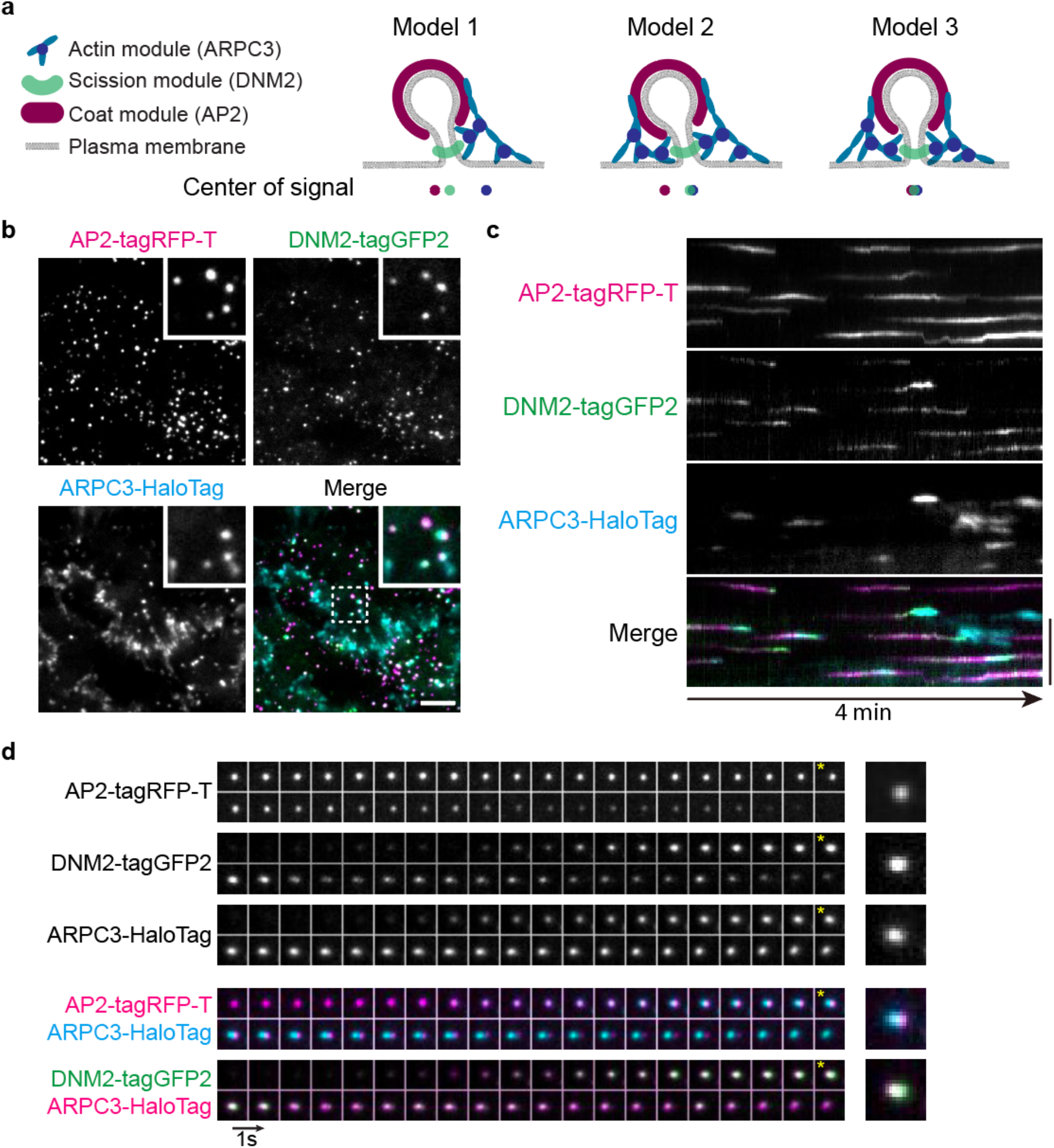
Triple-genome-edited iPS cells reveal dynamic actin organization at CME sites. **a**, Models of branched actin assembly at invaginating CME sites. Model 1: Asymmetric actin assembly at CME sites results in separated actin-coat and actin-neck signals. Model 2: Symmetric actin assembly at tilted CME sites results in separated actin-coat signals but overlapped actin-neck signals. Model 3: Symmetric actin assembly at perpendicularly invaginating CME sites will result overlapped actin, coat and neck signals. **b**, A representative single time frame image of a TIRF movie (Supplementary Video 2) of AP2M1-tagRFP-T (magenta), DNM2-tagGFP2 (green) and JF635 ligand^50^-conjugated ARPC3-HaloTag (cyan) in ADA cells. The highlighted region is boxed by a dashed line. Scale bar: 5µm. **c**, A representative kymograph of AP2M1-tagRFP-T (magenta), DNM2-tagGFP2 (green) and JF635 ligand-conjugated ARPC3-HaloTag (cyan) at CME sites in ADA cells. Scale bar: 5µm. **d**, Montage of a representative ARPC3 positive CME site in ADA cells. Individual channels and pair-wise merges are shown. *: Images from one frame before scission (maximum DNM2 intensity) are marked to show the displacement between the CME coat (AP2)-ARPC3 and CME neck (DNM2)-ARPC3. Size of field of view: 2µm × 2µm. Intervals: 1sec.

To analyze the intrinsic recruitment order and timing for up to three endocytic proteins at CME sites quantitatively and systematically, we developed an automated, high-throughput method that avoids bias because it does not involve manual selection of CME sites (see Materials and Methods). Briefly, AP2 tracks were identified using standard particle-tracking algorithms^30^. Novel filtering methods then extracted DNM2-positive events marked by one or more DNM2 burst. The AP2 and DNM2 tracks were decomposed into dynamic features describing the events’ position and brightness. These features were used for clustering via unsupervised machine learning, which enabled grouping of similarly-behaved tracks (Supplementary Fig. 3a and b). DNM2-positive events were refined by a detection scheme that determined the number of DNM2 peaks using various characteristics of a single DNM2-peak: the peak height, width, and minimum peak-to-peak distance (Supplementary Fig. 3c). Events with a single DNM2 peak were analyzed as described below. The method detects low signals from endogenously tagged CME proteins, such as the low-level recruitment of DNM2 at the early stages of CME, and accurately reveals the different CME stages (Extended Data Fig. 3d).

Next, the timing of actin network assembly at CME sites was determined using ARPC3 as a branched actin filament marker by analyzing over one thousand CME events. Although actin appearance early in CME has been reported^14^, determining the actin assembly timing is challenging because it is difficult to distinguish newly assembled branched actin at CME sites from the nearby cortical actin filaments or actin filaments attached to other vesicles or organelles. Also, whether actin functions during the early stage of CME has not yet been shown conclusively due to the potential side effects such as changes in membrane tension caused by actin inhibitors. Our endogenous ARPC3 tagging and large-scale computational analysis approach sidesteps these problems. We classified ARPC3 positive CME events into two groups: one group with ARPC3 appearance early in CME, and the other with late appearance in CME (Fig. 3a). We observed that in most of the events (N=1,385, 67.8%) a sharply increasing ARPC3 signal appears with similar timing to the rapid-recruitment phase of DNM2 concomitant with the U to Ω membrane shape transition. This timing is consistent with previously proposed role for actin in membrane invagination, as studies showed that actin inhibitors block the U to Ω endocytic membrane shape transition^6,14^. In some cases (N=657, 32.2%) we detected ARPC3 signals at early CME stages. To test whether random overlap between nearby actin structures and CME sites might be responsible for the apparent early actin recruitment, we generated a randomized data set by pairing ARPC3 images with AP2 and DNM2 images from an unrelated movie (Fig. 3b). In this data set, we detected early “assembly” of actin in the majority of ARPC3 positive CME events (N=17,282, 72.9%), and the intensity profiles of these events resembled the early-actin CME events we observed in the real data set (Fig. 3a, b). Therefore, we conclude that the presence of actin early in CME is very likely due to unrelated nearby actin structures overlapping with CME sites.

**Fig. 3:**
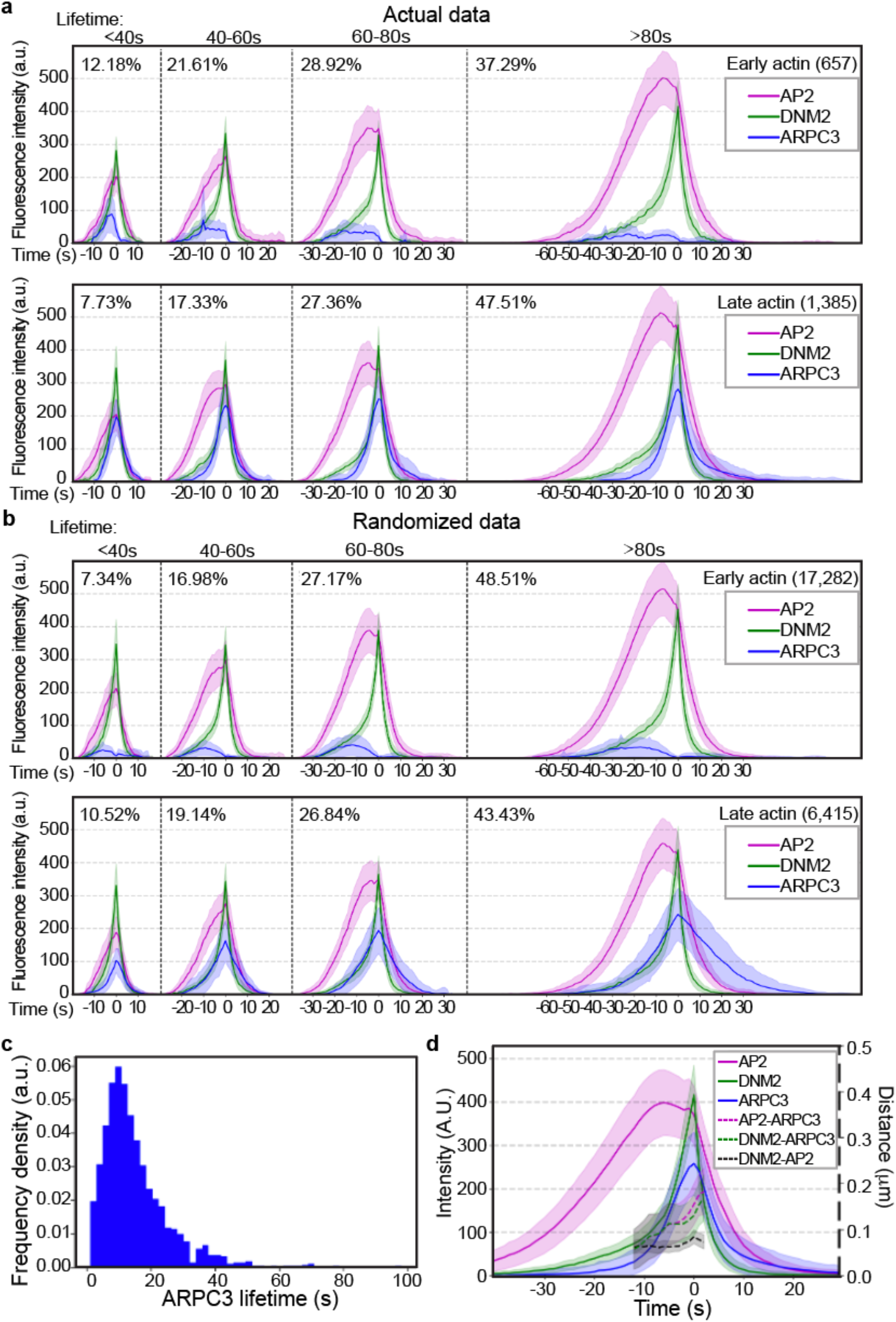
Computational analysis of ARPC3 positive CME sites reveals asymmetric actin network assembly at the late stage of CME. **a, b**, Averaged intensity vs time plots of cohorts of ARPC3 positive CME sites in ADA cells (**a**) and in the randomized data set (**b**). Events are grouped by the timing of ARPC3-labeled branched actin network recruitment (early: top, late: bottom), and then grouped into cohorts by the lifetimes of AP2 and aligned to the frames showing the maximum DNM2 intensity (time = 0s). Total number of CME sites in each group is shown in parentheses. Percentage of the number of the CME sites in each cohort is shown next to the plot. **c**, Histogram of ARPC3-mediated actin network assembly duration. The assembly duration is measured from the first frame of the ARPC3 signal to the presumed scission time (the peak of DNM2 signal). **d**, Averaged intensity (solid lines) and distance (dashed lines) vs time plots of ARPC3 positive CME sites in ADA cells. Events are aligned to the frames showing the maximum DNM2 intensity (time = 0s). Distance between centers of two signals are shown from -10s to 3s when DNM2 and ARPC3 signals are relatively high. N=1,385. **a, b, d**, Error bar: ¼ standard deviation.

Our live-cell analysis allowed the timing of branched actin network assembly to be compared to the scission timing, and the spatial offset between the clathrin coat and the associated actin network to be determined. Super-resolution imaging of yeast CME sites suggested that actin and actin nucleators localize symmetrically in a ring around CME sites, and computational modeling suggested that an asymmetric actin arrangement would not provide sufficient force for the membrane invagination during yeast CME^13^. In contrast, in mammalian cells, which require less actin force production during CME, imaging of fixed cells suggested that actin structures associate adjacent to apparent flat clathrin coats. However, these studies proposed that at the later CME stages the actin structures become larger and more symmetric to provide sufficient force for membrane deformation and scission^14,15^. Surprisingly, in our live cell studies designed to highlight sites of new actin assembly, we observed off-centered branched actin networks at CME sites throughout even the latest CME stages (Fig. 2d). Furthermore, most ARPC3-positive CME sites accomplish scission within 30s from the initiation of ARPC3 recruitment (Fig. 3c). The actin networks were off center from the coat and neck signals by approximately 150nm at the time of vesicle scission (Fig. 3d). Given the temporal separation between channel acquisition and the movement of AP2 spots, the separation between channels can be attributed in part to an imaging artifact. Therefore, when we measured the average movement of AP2 spots leading up to scission as a control, we found that over 95% of the events had AP2-ARPC3 separations that exceed the frame-to-frame motility of AP2 (Supplementary Fig 4). Also, the uncertainties measured by a standard deviation, when measuring the fitted position of AP2, range up to 40 nm. Therefore, we include the AP2-DNM2 separation as a basis for comparison to the AP2-ARPC3 and DNM2-ARPC3 separations (Fig 3d). These results further support our conclusion that branched actin networks assemble asymmetrically at CME sites through the time of scission (Fig 2d). This observation is consistent with the observation that ring-shaped actin structures at clathrin coats were rarely observed in the high-resolution, live-cell imaging in a previous study^31^. In total, these live-cell data suggest that in mammalian cells, asymmetric actin network assembly can provide enough force to assist membrane deformation and scission during the late stages of CME.

### Asymmetric branched actin networks facilitate CME at stalled sites

To gain additional insights into the function of this asymmetric actin network assembly, we quantitatively compared kinetics of CME events with or without ARPC3 recruitment. We observed that about 30% of CME events are completed in the absence of detectable actin assembly, which is consistent with the hypothesis that in mammalian cells actin assembly is required for CME only under relatively high membrane tension, which can vary regionally within cells^6,7,9^. Consistent with the possibility that increased membrane tension stalls membrane deformation during CME^4,8,9,32–34^, CME lifetimes were markedly longer for ARPC3 positive events compared to the ARPC3 negative events (Fig. 4a). In addition, when the AP2M1 intensity vs time profiles were compared between ARPC3 positive and negative CME sites, a plateau, which lasts for approximately 10 seconds, was observed for the ARPC3 positive events (Fig. 4b). Based on these observations and previous experimental and computational modeling data^4,6,7^, we propose that this plateau in branched actin-positive CME events represents stalled membrane bending due to an unfavorable local membrane environment, such as higher membrane tension^4,32,33^.

**Fig. 4:**
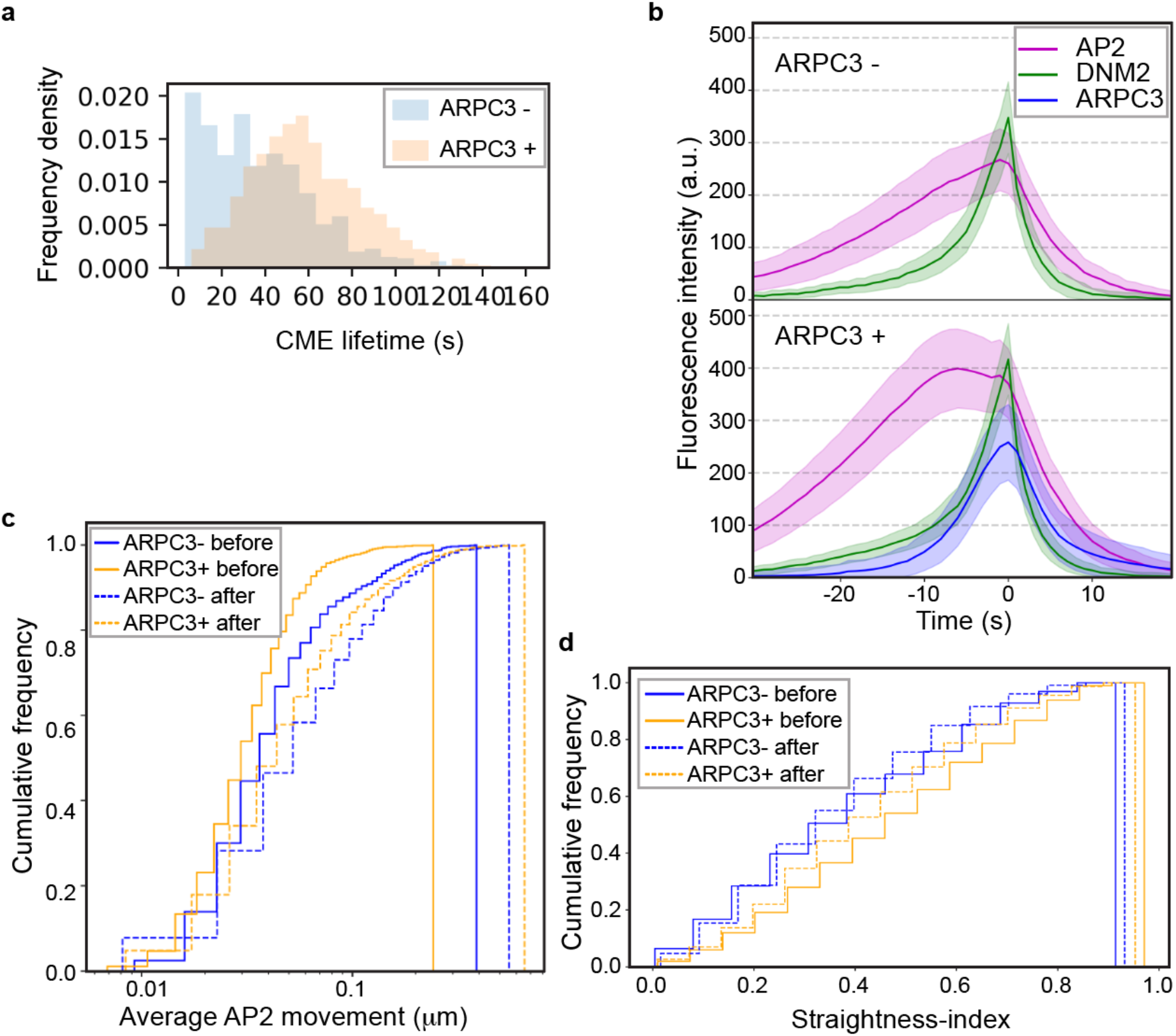
Actin positive CME sites show distinct dynamics. **a**, Histograms of ARPC3 negative (blue) and positive (orange) CME lifetimes. CME lifetime is measured from the first frame of the AP2 signal to the presumed scission time (the peak of DNM2 signal). ARPC3 positive CME events have longer lifetimes. **b**, Averaged intensity vs time plots of ARPC3 negative (top) and positive (bottom) CME sites in ADA cells. Events were aligned to the frames showing the maximum DNM2 intensity. Error bar: ¼ standard deviation. **c**, Lateral motility of ARPC3 negative (blue) and positive (yellow) CME sites before (solid line) and after (dashed line) vesicle scission. ARPC3 positive CME sites move slower than ARPC3 negative ones. **d**, Straightness-index of ARPC3 negative (blue) and positive (yellow) CME sites before (solid line) and after (dashed line) scission. The straightness-index is defined by the ratio between the sum of frame-to-frame distances to the end-to-end distance of a single event’s trajectory, where a perfectly straight-lined trajectory would have an index of 1. APRC3 positive CME sites move with a straighter trajectory. **a-d**, ARPC3 -: N=840, ARPC3 +: N=1,385.

We next tested the hypothesis that the asymmetric actin network might affect the lateral movements of endocytic coats on the plasma membrane. Interestingly, the ARPC3 positive CME sites showed significantly slower, but more directional lateral movement before the scission compared to the ARPC3 negative CME sites (Fig. 4c, d). After scission both ARPC3 positive and negative vesicles showed fast, apparently random movements (Fig. 4c, d). These data suggest that the asymmetric actin can stabilize the forming endocytic coat while pushing it in the plane of the plasma membrane with a lateral directional force.

### N-WASP is recruited asymmetrically to the stalled CME sites

To further explore how the asymmetrical assembly of actin networks at CME sites is regulated, we endogenously tagged N-WASP, an actin nucleation promoting factor (NPF) that plays roles in CME, in AP2M1-tagRFP-T/ DNM2-tagGFP2 genome-edited iPSCs (hereafter referred to as ADW cells, Fig. 5a and Supplementary Fig. 5a). Quantitative imaging of budding yeasts demonstrated that initiation of productive actin assembly at CME sites requires the accumulation of yeast WASP or WIP to a certain amount^35^. In our genome-edited iPS cells, we observed that N-WASP is recruited asymmetrically to CME sites mostly at the late stage of CME (Fig. 5b, c and Supplementary Fig. 5b, c). Longer lifetimes and a plateau in the AP2 intensity vs time plot were observed specifically in the N-WASP positive CME events (Fig. 5d, e), similar to the ARPC3 positive events (Fig 4 a, b). These data indicate that asymmetric NPF recruitment underlies the asymmetric architecture of branched actin networks at CME sites.

**Fig. 5:**
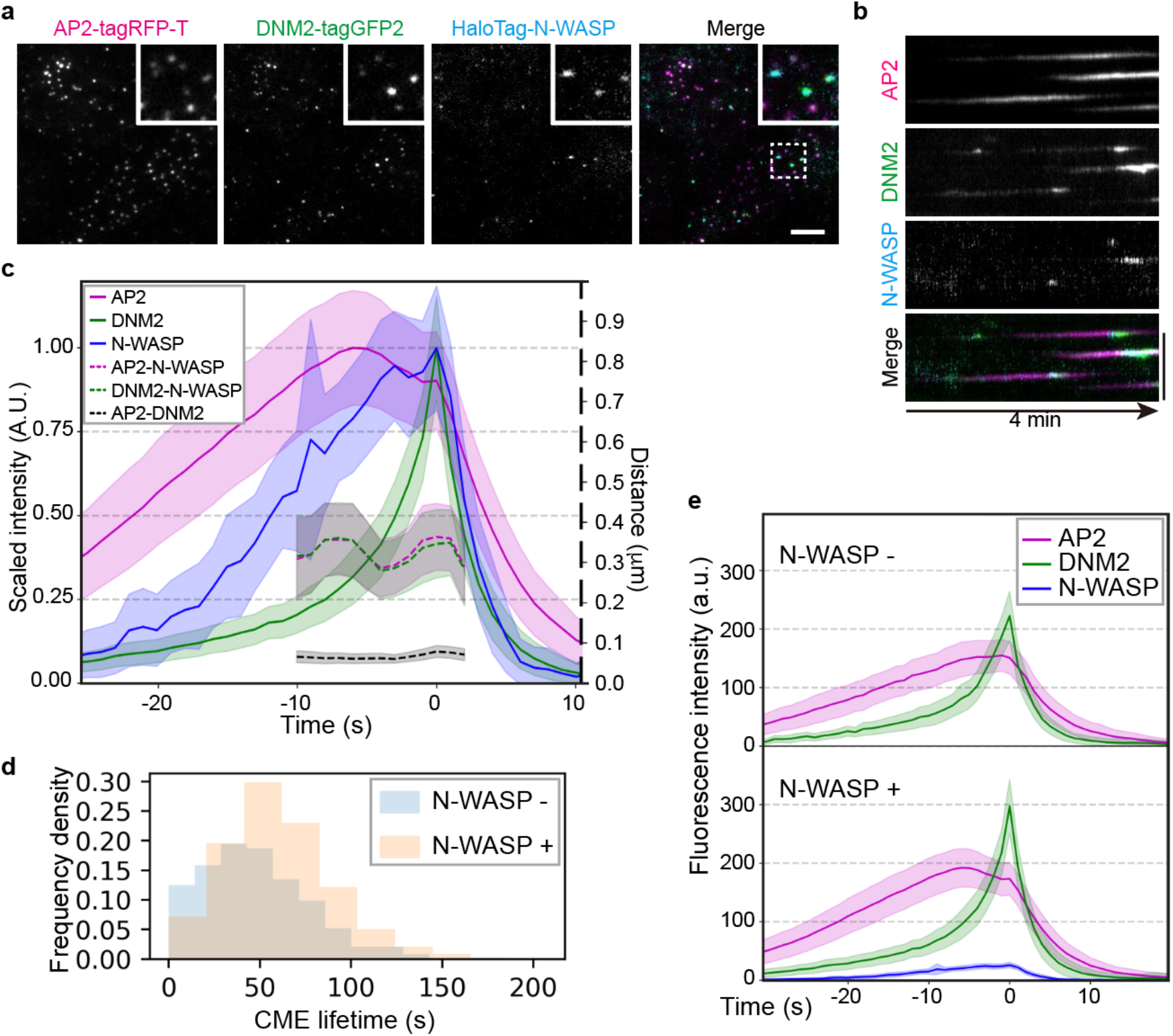
Asymmetric N-WASP recruitment to stalled CME sites. **a**, A representative single time frame image of a TIRF movie (Supplementary Video 4) of AP2M1-tagRFP-T (magenta), DNM2-tagGFP2 (green) and JF635 ligand-conjugated HaloTag-N-WASP (cyan) in ADW cells. The highlighted region is boxed by a dashed line. Scale bar: 5µm. **b**, A representative kymograph of CME sites in ADW cells over a 4 min movie. Scale bar: 5µm. **c**, Averaged intensity (solid line) and distance (dashed line) vs time plots of N-WASP positive CME sites in ADW cells. Events are aligned to the frames showing the maximum DNM2 intensity. Intensity is scaled to 1 at peaks for each channel. Error bar: ¼ standard deviation. **d**, N-WASP positive CME events have longer lifetimes. **e**, Intensity vs time plots of averaged N-WASP negative (top) and positive (bottom) CME sites in ADW cells. Events were aligned to the frames showing the maximum DNM2 intensity. Error bar: ¼ standard deviation. **c-d**, N-WASP negative CME sites: N=385, N-WASP positive CME sites: N=1,381

## Discussion

Using large-scale, comprehensive analysis of thousands of CME sites in unperturbed live cells, our study demonstrates that in mammalian cells coat assembly dynamics predict which sites will assemble actin, and show that at apparently stalled sites, actin assembles asymmetrically to facilitate successful vesicle formation.

Based on the data presented here, we propose an updated model for actin assembly at mammalian CME sites in which, beyond global tension-dependent changes in requirement for actin assembly, highly localized differences give rise to heterogeneity even within the same patch of plasma membrane in the same cell (Fig. 6): (1) Where the local membrane tension is lower (Fig. 6 upper scenario), the membrane can undergo flat-U-Ω shape transitions without actin assembly in a relatively short time. When the coat grows large enough to form a Ω-shaped bud, sufficient dynamin can be recruited to perform scission, and there is little delay between coat expansion and scission; (2) Where the local conditions are not favorable, presumably under high membrane tension and possibly other impediments, the coat protein-membrane interaction does not generate sufficient force to curve the membrane (Fig. 6 lower scenario). Here, extra force generation from actin assembly is required^4,6^. Asymmetric N-WASP recruitment activates actin nucleation mostly at one side of the clathrin coat, generating an asymmetric force that pulls the membrane into the cell with a similar action to a bottle cap opener. We speculate that this asymmetrical force contributes to asymmetric membrane deformation at endocytic sites observed by high-speed atomic force microscopy^36^ and may act with dynamin^37^ to twist the clathrin pit to promote scission at the neck. CME events with associated actin assembly have longer lifetimes, likely due to a delay between coat expansion and scission, requiring adaptive recruitment of actin regulators followed by actin network assembly and membrane remodeling. This result reinforces the conclusion from previous studies^6–9^ that increased membrane tension enhances the requirement for actin assembly during CME, but also establishes that site-to-site heterogeneity in actin dependence and involvement can be observed without manipulating membrane tension.

**Fig. 6:**
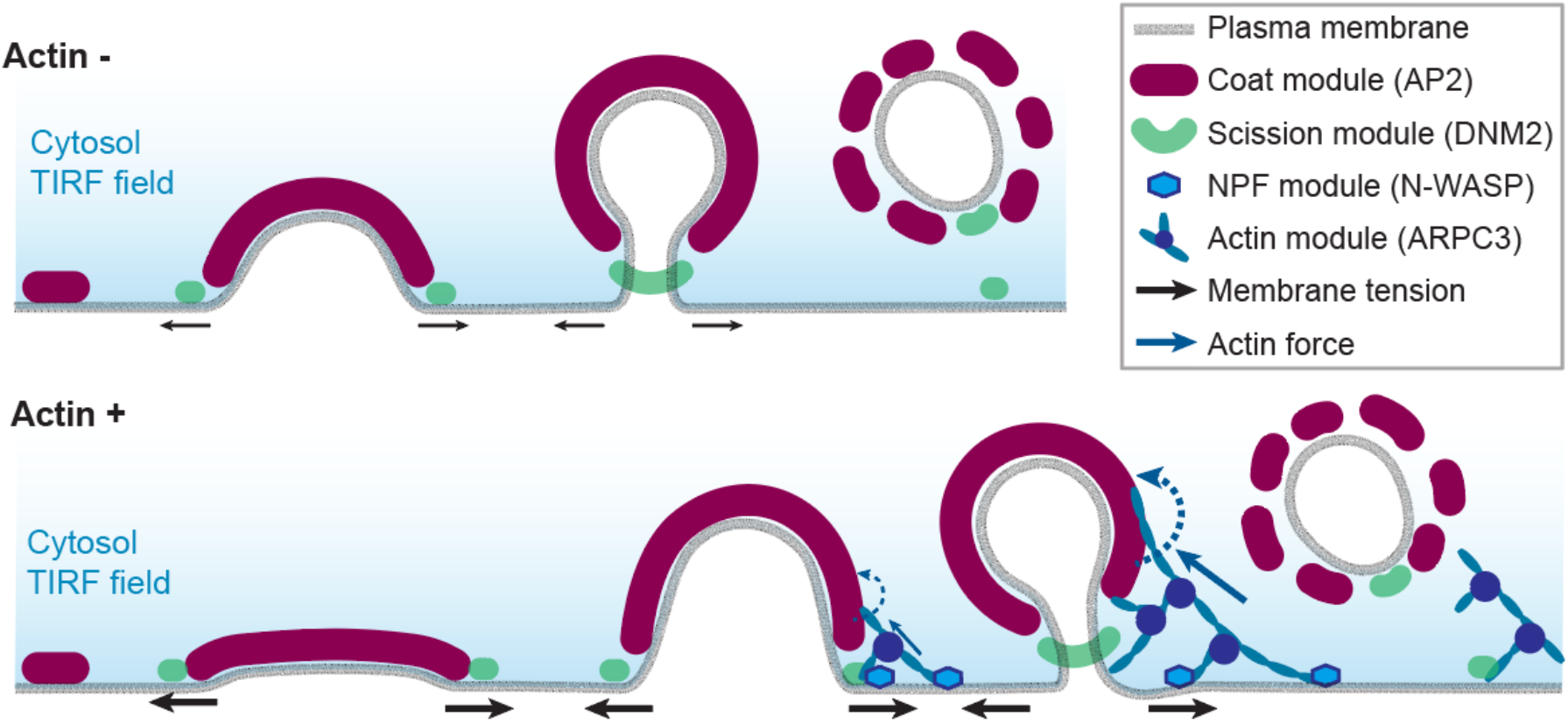
An updated schematic model of actin-negative and actin-positive clathrin-coated pits in human cells. Actin assembly is induced at stalled CME sites, where asymmetric forces pull, bend and possibly twist the plasma membrane against membrane tension to drive membrane invagination and vesicle scission.

Future computational modeling studies of how asymmetric actin network assembly provides forces for vesicle formation and membrane remodeling will deepen our understanding of actin’s functions in a host of actin-mediated processes.

Our model provides further insights into the basis for inconsistent effects of actin drugs on CME^6,14,18,38–43^. Actin plays crucial roles in membrane shaping, cell adhesion, and membrane tension. Global disruption of actin dynamics is expected to dramatically change membrane tension and the available pool of actin and associated proteins and therefore to have both direct and indirect effects on CME. Here, we focused on in-depth analysis of the unperturbed process and detected preference for actin assembly at stalled CME events.

The results presented here may prove relevant to the constant coat area vs constant coat curvature debate for how the clathrin coat assembles and develops curvature^32,44–48^. In the constant area model, flat clathrin coats grow close to their final size before curvature develops as a vesicle forms. In the constant curvature model, clathrin coats grow with a fixed curvature. Our observations suggest that coat expansion and curvature generation may be regulated via distinct mechanisms, with different actin requirements. At actin-positive CME sites, actin assembles primarily at the late stage of CME when coat assembly is mostly complete (Fig. 3 and Supplementary Fig. 6a). If the constant curvature model holds, actin should only be associated with clathrin coats with highly curved dome and spherical shapes (Supplementary Fig. 6b). However, actin associated with flat or shallow clathrin coats has been observed in multiple studies^7,15^, which supports the constant area model at actin-positive CME sites (Supplementary Fig. 6c). On the other hand, mathematical modeling predicts that at actin-negative CME sites, where coat and other curvature-promoting proteins provide sufficient force to bend the membrane^23^, the membrane is smoothly and continuously shaped by these proteins into a budded morphology as the coat area increases^4^, which follows the constant curvature model. Perhaps the constant area model applies primarily for actin-negative sites and the constant area model applies primarily for actin-positive sites, which we have shown here are mostly stalled CME sites. We suggest that in future studies the constant coat area and constant coat curvature models be tested at individual CME events to test the possibility that both mechanisms operate in the same cell.

## Acknowledgments

MJ was funded by American Heart Association Postdoctoral Fellowship (18POST34000029). DGD was funded by NIH MIRA grant R35GM118149. KX is a Chan Zuckerberg Biohub investigator and acknowledges support from NIH (DP2GM132681). SU was funded by Philomathia Foundation and the Chan Zuckerberg Initiative Imaging Scientist program. The authors would like to thank Dr. Yidi Sun and Dr. Matthew Akamatsu for insightful comments on the manuscript; the Conklin Lab at UCSF for providing WTC10 human iPSC line; the Lavis Lab at Janelia Research Campus for providing JF635 HaloTag ligand; Dr. Sun Hae Hong for generating the AP2-tagRFP-T iPSC cell line; the UC Berkeley QB3 MacroLab for purified S. pyogenes NLS-Cas9; the UC Berkeley Cancer Research Laboratory Flow Cytometry Facility for iPSC sorting.

## Data availability

The raw live-cell imaging data (TIRF) can be found at https://github.com/DrubinBarnes/Jin_Shirazinejad_et_al_branched_actin_manuscript. All other raw data are available from the corresponding author upon request.

## Code availability

The Jupyter Notebooks used for live-cell imaging analysis can be found at https://github.com/DrubinBarnes/Jin_Shirazinejad_et_al_branched_actin_manuscript.

## METHODS

### Cell culture

The WTC10 hiPSC line was obtained from the Bruce Conklin Lab at UCSF. hiPSCs were cultured on Matrigel (hESC-Qualified Matrix, Corning) in StemFlex medium (Thermo Fisher) with Penicillin/ Streptomycin in 37°C, 5% CO2. Cultures were passaged with Gentle Cell Dissociation reagent (StemCell Technologies, Cat#: 100-0485) twice every week.

### Genome-editing

The AP2M1 gene was edited in WTC10 hiPSCs as previously described using TALENs targeting exon 7 of the AP2M1 gene^49^. Both alleles of AP2M1 were tagged with tagRFP-T. The Cas9-crRNAtracrRNA complex electroporation method was used sequentially to edit DNM2 and ARPC3 gene in AP2M1-tagRFP-T genome-edited hiPSCs, as previously described^12,18^. The same method was used to edit the WASL gene in AP2M1-tagRFP-T/DNM2-tagGFP2 genome edited hiPSCs. *S. pyogenes* NLS-Cas9 was purified in the University of California Berkeley QB3 MacroLab. TracrRNA and crRNA that target CCTGCTCGACTAGGCCTCGA (DNM2), CCTGGACAGTGAAGGGAGCC (ARPC3) and AGCTCATGGTTTCGCCGGCG (WASL), were purchased from IDT. Gibson assembly (New England Biolabs) was used to construct donor plasmids containing DNM2 5’ homology-ggtaccagtggcggaagc-tagGFP2-DNM2 3’ homology, ARPC3 5’ homology-ggatccggtaccagcgatccaccggtcgccacc-HaloTag-ARPC3 3’ homology, and WASL 5’ homology-HaloTag-agcgatccaccggtcgccaccggatcc-WASL 3’ homology sequences, respectively. Three days after electroporation (Lonza, Cat#: VPH-5012) of the Cas9-crRNA-tracrRNA complex and donor plasmid, the tagGFP2 or HaloTag positive cells were single cell sorted using a BD Bioscience Influx sorter (BD Bioscience) into Matrigel-coated 96-well plates. Clones were confirmed by PCR and Sanger sequencing of the genomic DNA locus around the insertion site. Both alleles of DNM2 and ARPC3 were tagged with tagGFP2 and HaloTag, respectively, and one allele of WASL was tagged with HaloTag in the hiPSC lines used in this study.

### Western blotting

Cells were dissociated from the well using Gentle Cell Dissociation reagent (StemCell Technologies, Cat#: 100-0485). Total proteins were extracted by adding 1ml of cold 10% TCA to the cell pellets, incubated on ice for 30min, and spun down by centrifuging at 4 °C, 12000rpm for 10min. Protein pellets were dissolved in loading buffer (50 mM HEPES, pH 7.4, 150 mM NaCl, 1 mM MgCl2, 5% BME, 5mM DTT and protease inhibitor) and loaded onto an acrylamide gel for SDS-PAGE and transferred to nitrocellulose membranes for immunoblotting. Blots were incubated overnight at 4°C with primary antibodies targeting Tag(CGY)FP (1:2000 dilution in 1% milk, Evrogen, Cat#: AB121), HaloTag (1:1000 dilution in 0.5% milk, Promega, Cat#: G9211), GAPDH (1:100,000 dilution in 0.5% milk, Proteintech, Cat#: 10494-1-AP), respectively, and subsequently incubated in the dark at room temperature for 1hr with secondary antibodies.

### TIRF live-cell imaging

Two days before imaging, hiPSCs were seeded onto Matrigel-coated 4-well chambered cover glasses (Cellvis). Halotag was labeled by JF635-HaloTag ligand^50^. Cells were incubated in StemFlex medium with 100 mM JF635-HaloTag for 45min and the unbound ligands were washed away by three washes with 5 min incubation in prewarmed StemFlex medium. Cells were imaged on a Nikon Ti-2 inverted microscope fitted with TIRF optics and a sCMOS camera (Hamamatsu). Cells were maintained at 37 °C with a stage top incubator (OKO Lab) in StemFlex medium with 10mM HEPES. Images were acquired with Nikon Elements. Channels were acquired sequentially at a 1 sec interval and 300ms exposure time over 4 minutes.

### TIRF image processing

Four generalized processing steps were applied identify of clathrin-coated pits with single DNM2 peaks: track feature abstraction, feature dimensionality reduction, event clustering, and DNM2-peak detection. First, tracks that are defined by fitted positions and intensities for single events were generated using cmeAnalysis^30^. Then, AP2 and DNM2 tracks were decomposed into dynamic features describing the dynamics of the events’ position and brightness. The mapping of each track to discrete features was done to generalize the dynamics of tracked events into a set of interpretable coordinates. These features were clustered after feature scaling to normal distributions, dimensionality reduction with principal component analysis, and Gaussian mixture modeling. DNM2-positive events represented a distinct cluster of tracks that had detectable DNM2 throughout the event, were long lived, and were below the threshold of motility expected for transient, non-CME-derived clathrin-coated vesicle “visitors” at the TIRF field. Single DNM2-peak events were found by searching over a range of values set for the minimum DNM2 peak height, width, and peak-to-peak temporal distance. After finding single-peaked events in a fixed peak-parameter combination, the lifetime distribution of single peak events’ lifetimes were fit to the expected underlying distribution, a Rayleigh distribution^51^, where the best-fitting parameter combination was chosen to identify single-peaked events. Single DNM2-peaked events were kept as CME sites for the remainder of the analysis. All code associated with this analysis, generating Figures 3-5, and a detailed step-by-step protocol, are available at https://github.com/DrubinBarnes/Jin_Shirazinejad_et_al_branched_actin_manuscript.

### Two-color 3D STORM imaging

12 mm round coverslips were sonicated in distilled water and sterilized for 20 min in 70% ethanol, air-dried and coated with Matrigel in 24-well plates. Cells were seeded onto Matrigel-coated coverslips two days before fixation. For clathrin and actin two-color imaging, cells were fixed first for 1 min in 0.3% (v/v) glutaraldehyde (GA) solution containing 0.25% (v/v) Triton in cytoskeleton buffer (CB: 10mM MES, 150mM NaCl, 5mM EGTA, 5mM Glucose, 5mM MgCl2, 0.005% NaN3, pH 6.1) and then immediately fixed for 20 min in 2% (v/v) GA solution in CB. Both solutions were prepared fresh from a 10% GA stock (Electron Microscopy Science, cat #16120). After fixation, samples were incubated twice for 5 min in freshly prepared 0.1% (w/v) NaBH4 in PBS. For clathrin and ARPC3-HaloTag imaging, cells were fixed for 20 min in 4% (v/v) PFA (Electron Microscopy Sciences, Cat#: 15710) in CB. Subsequently, both types of samples were washed 3 times for 10 min in PBS. Samples were then blocked for 20 min in blocking buffer [3% (w/v) BSA and 0.1% (w/v) Saponin in PBS]. Clathrin light chain (Invitrogen, Cat#: MA5-11860, 1:200 dilution) and Halotag (Promega, Cat#: G9281, 1:200 dilution) antibodies were used in blocking solution. Primary antibody immunostaining was performed overnight at 4°C. On the next day, samples were washed three times in washing buffer (0.1x blocking buffer in PBS) for 10 min. Samples were incubated with secondary antibody in blocking buffer for 30 min at room temperature in the dark and were washed three times for 10 min in washing buffer, and then three times for 10 min in PBS. Homemade mouse secondary antibody-CF680 (1:50) was used to stain clathrin and actin samples. Commercial mouse secondary antibody-AF647 (ThermoFisher, cat#A32787; 1:400) and homemade rabbit secondary antibody-CF680 (1:50) were used to stain the clathrin and ARPC3-HaloTag. Clathrin and actin samples were then stained with 0.5µM Phalloidin-AF647 (Fisher Scientific, Cat#: A22287) in PBS and kept at room temperature in the dark for 2 hours. Samples were washed three times with PBS before STORM imaging.

STORM imaging was performed as previously described on a homebuilt STORM setup^7,52^. Samples labeled by AF647 and CF680 were excited by an 647nm laser. The emission of both AF647 and CF680 was then split into two light paths as two channels using a dichroic mirror (Chroma, cat#T685lpxr), and each channel was projected onto one-half of an EMCCD camera (Andor iXon Ultra 897). Color assignment of each localization was based on its intensity in the two channels. A cylindrical lens was inserted into the transmitted channel to acquire 3D localization^28^. 3D position of each localization was determined from the ellipticity of each point spread function.

---

**Supplementary Fig 1:**
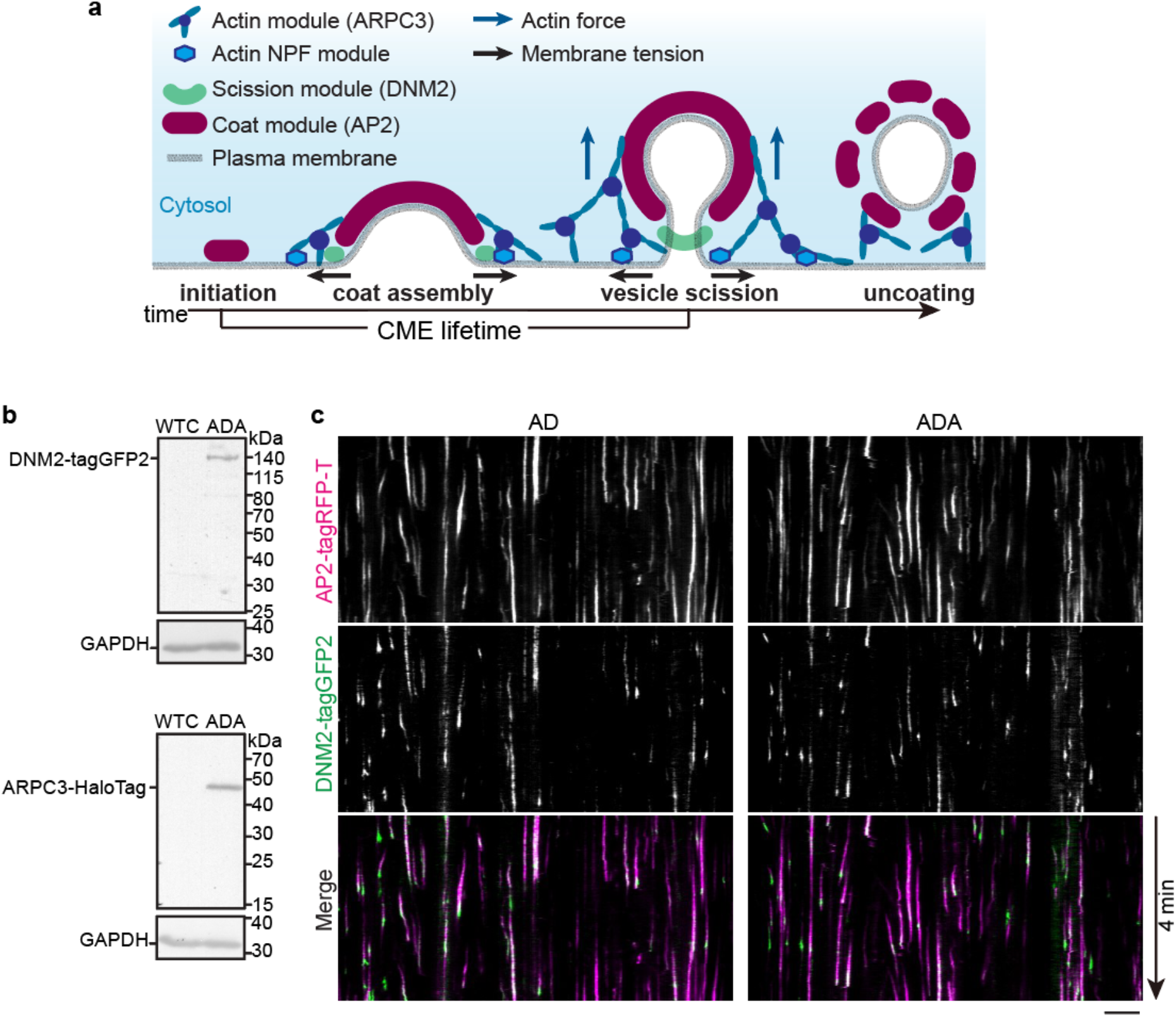
Genome-edited iPSCs show dynamic CME sites. **a**, Schematic model of CME. Mammalian CME proteins can be grouped into several modules, including the coat, WASP and Myosin / actin nucleation promoting factor (NPF), actin and scission modules^5^. Actin networks provide pulling forces to invaginate the membrane against membrane tension^4,6,9,12^. **b**, Immunoblot analysis of cell extracts from the WT (WTC) and genome-edited (AP2M1-tagRFP-T/DNM2-tagGFP2/ARPC3-HaloTag; ADA) human iPSCs. The labeled proteins were detected with tag(CGY)FP, HaloTag, and GAPDH (loading control) antisera respectively. **c**, Kymograph of representative CME sites of double-edited (AP2M1-tagRFP-T/DNM2-tagGFP2; AD) and triple-edited (AP2M1-tagRFP-T/DNM2-tagGFP2/ARPC3-HaloTag; ADA) cells.

**Supplementary Fig 2.**
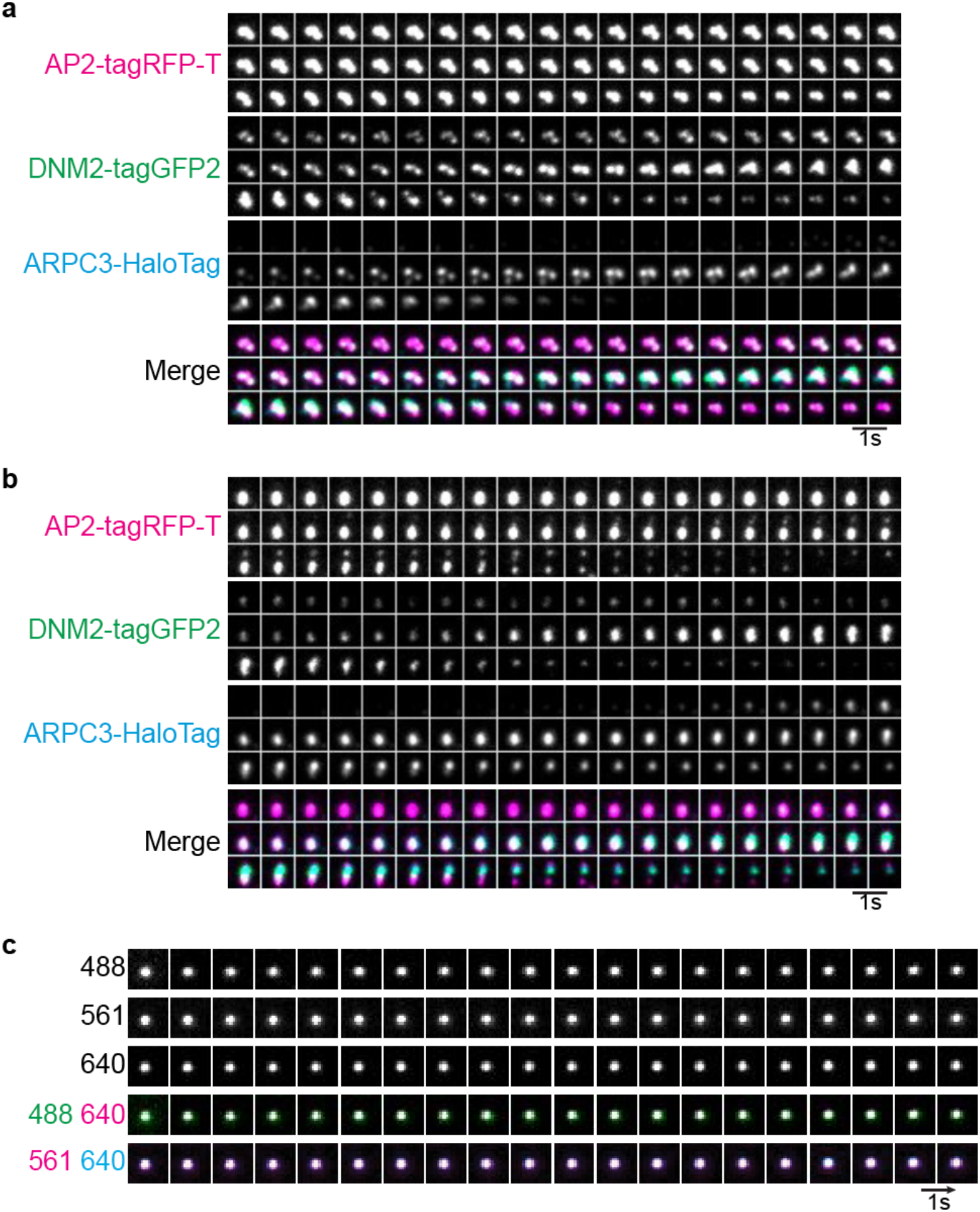
Actin assembles at different types of CME sites. **a**, Montage of a representative ARPC3 positive CME plaque from a TIRF movie of triple-edited (AP2M1-tagRFP-T/DNM2-tagGFP2/ARPC3-HaloTag; ADA) human iPSCs (Supplementary Video 2). **b**, Montage of a representative ARPC3 positive splitting CME site from a TIRF movie of triple-edited (AP2M1-tagRFP-T/DNM2-tagGFP2/ARPC3-HaloTag; ADA) human iPSCs (Supplementary Video 2). **c**, Montage from a TIRF movie of a multi-fluorescence bead (Supplementary Video 3). Size of field of view: 2µm × 2µm. Intervals: 1sec.

**Supplementary Fig 3.**
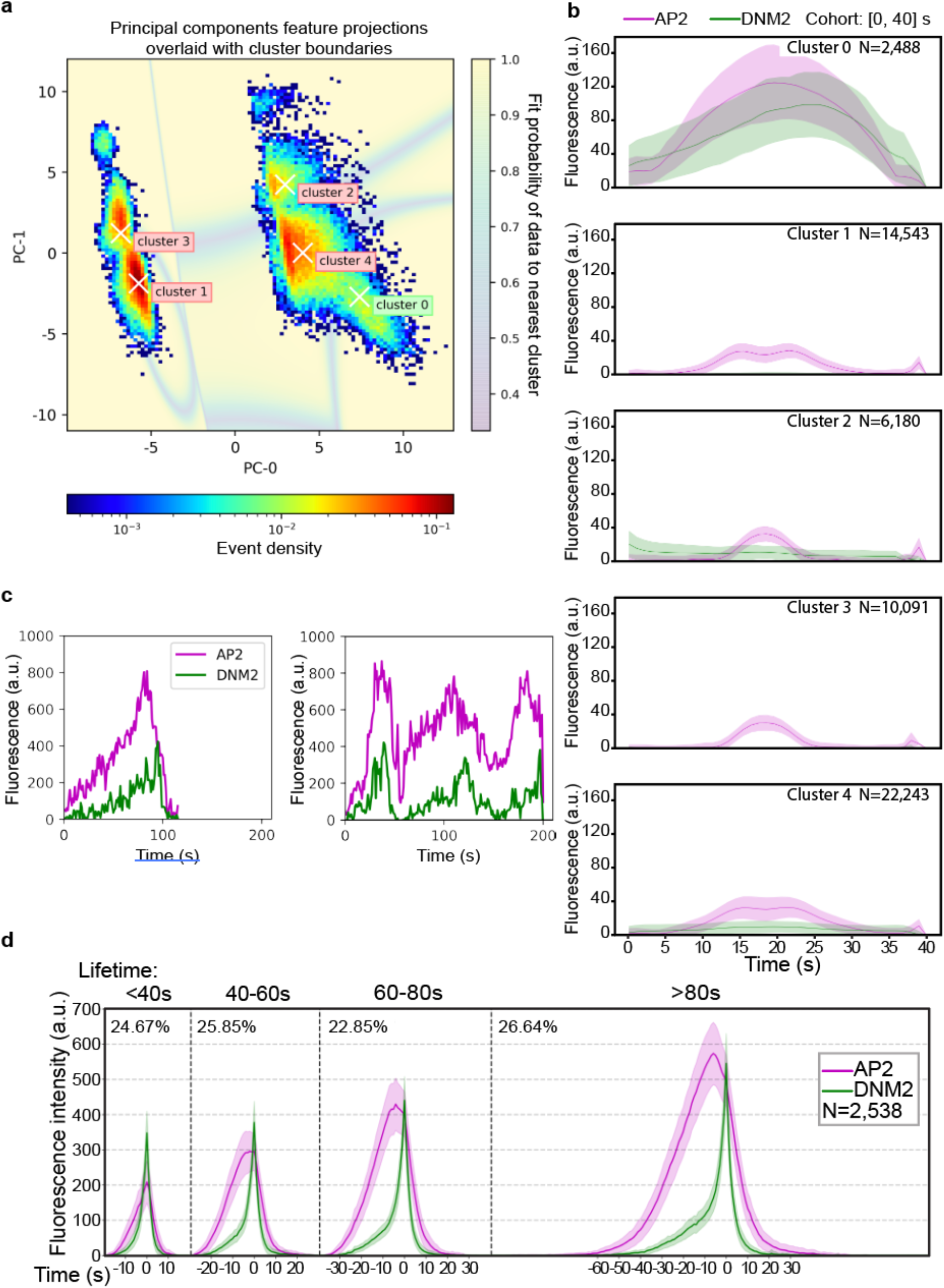
Filtering methods for selection of CME sites. **a**, 2-D histogram of the first two principal components (PCs) of AP2 and DNM2 dynamic features. The shaded underlay represents simulated data points in principal component space and their individual probabilities of belonging to the nearest cluster center. Cluster 0 shows data points in the DNM2-positive cluster. **b**, Cohort plots of the shortest AP2 events (<40 seconds) from each cluster. Cluster 0 represents DNM2-positive events where a strong DNM2 signal is detected. **c**, DNM2-positive events are sorted by the number of DNM2 peaks using a peak-detection scheme. Representative intensity vs time plots of a single-peaked event (left) and a multi-peaked event (right). **d**, Single-peaked DNM2 events, hereon named CME sites, are grouped into lifetime cohorts and aligned to the peak of the DNM2 channel.

**Supplementary Fig 4:**
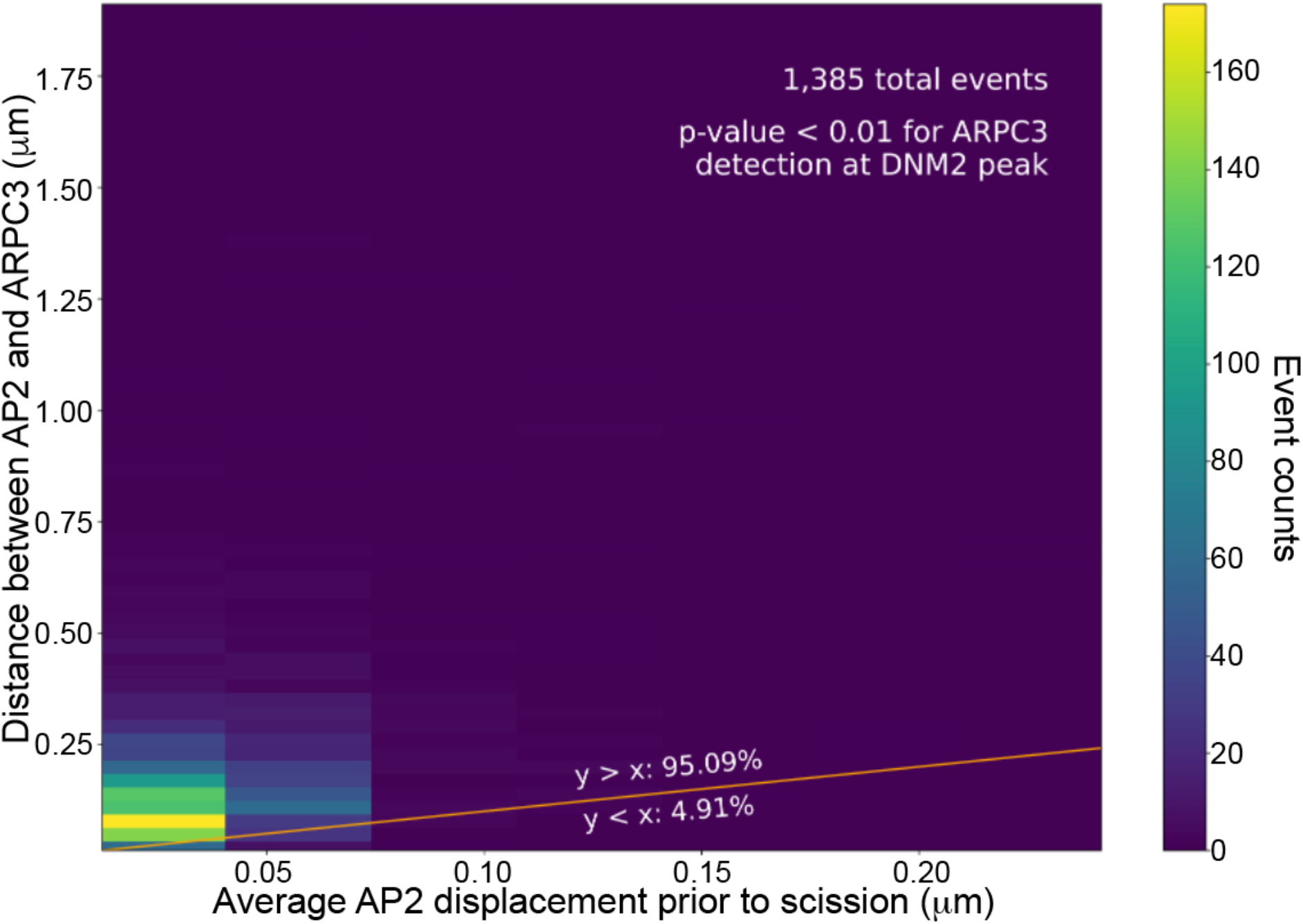
AP2-ARPC3 separation is not due to imaging artifacts. A heat map graph of distance between AP2 and ARPC3 signals before scission, and average AP2 frame to frame displacement within 6 seconds before scission. Over 95% of the CME events present larger AP2-ARPC3 separation than AP2 displacement. N= 1,385.

**Supplementary Fig 5:**
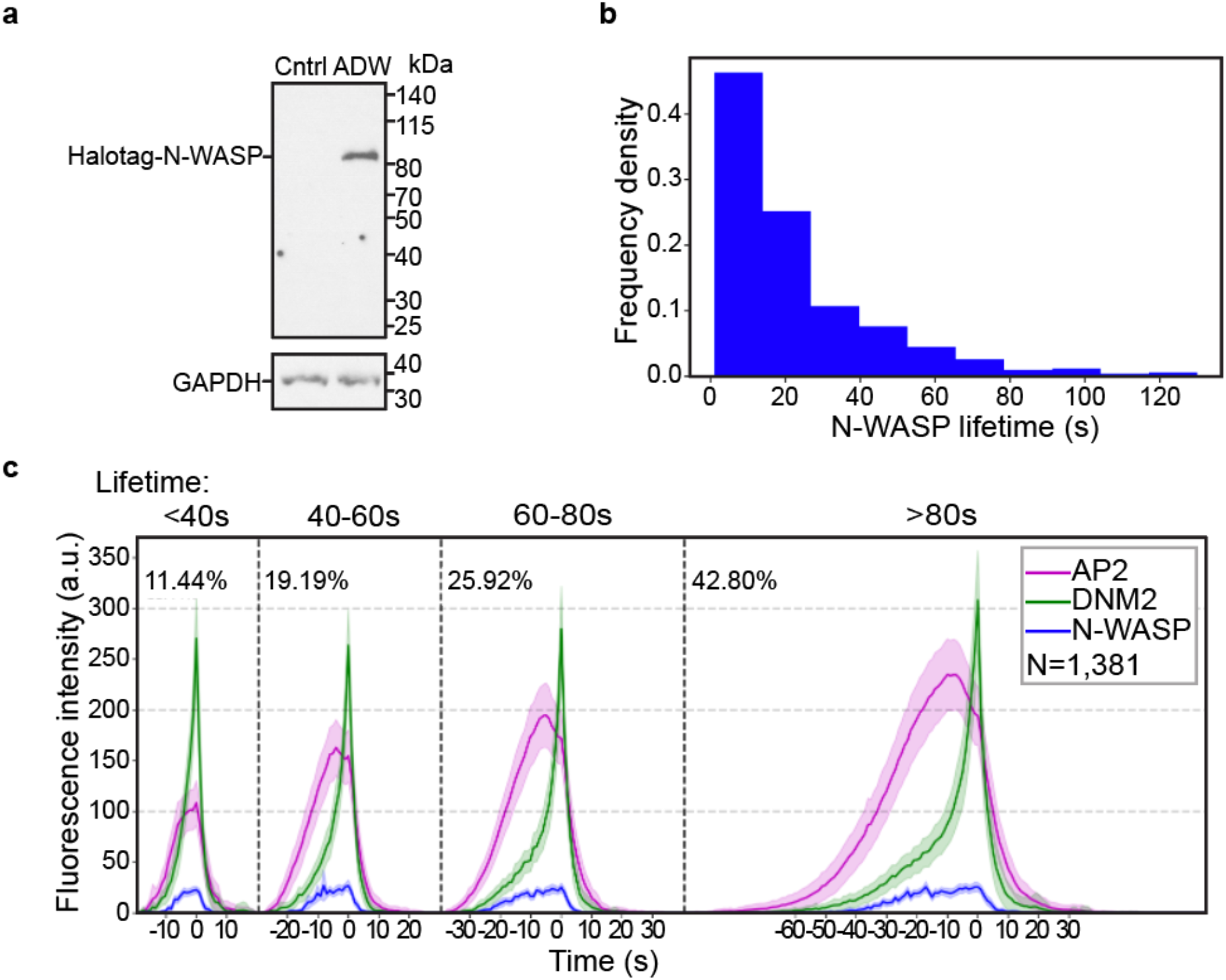
Dynamics of N-WASP at CME sites. **a**, Immunoblot analysis of cell extracts from the control and genome-edited (AP2M1-tagRFP-T/DNM2-tagGFP2/HaloTag-WASL; ADW) human iPSCs. The labeled proteins were detected with HaloTag and GAPDH (loading control) antisera respectively. **b**, Histogram of N-WASP lifetime at CME sites. The lifetime is measured from the first frame of the N-WASP signal to the presumed scission time (the peak of DNM2 signal). **c**, Intensity vs time plots of cohorts of N-WASP positive CME sites in ADW cells. Events are grouped into cohorts by the lifetimes of AP2 and aligned to the frames showing the maximum DNM2 intensity. N=1,381. Error bar: ¼ standard deviation.

**Supplementary Fig 6:**
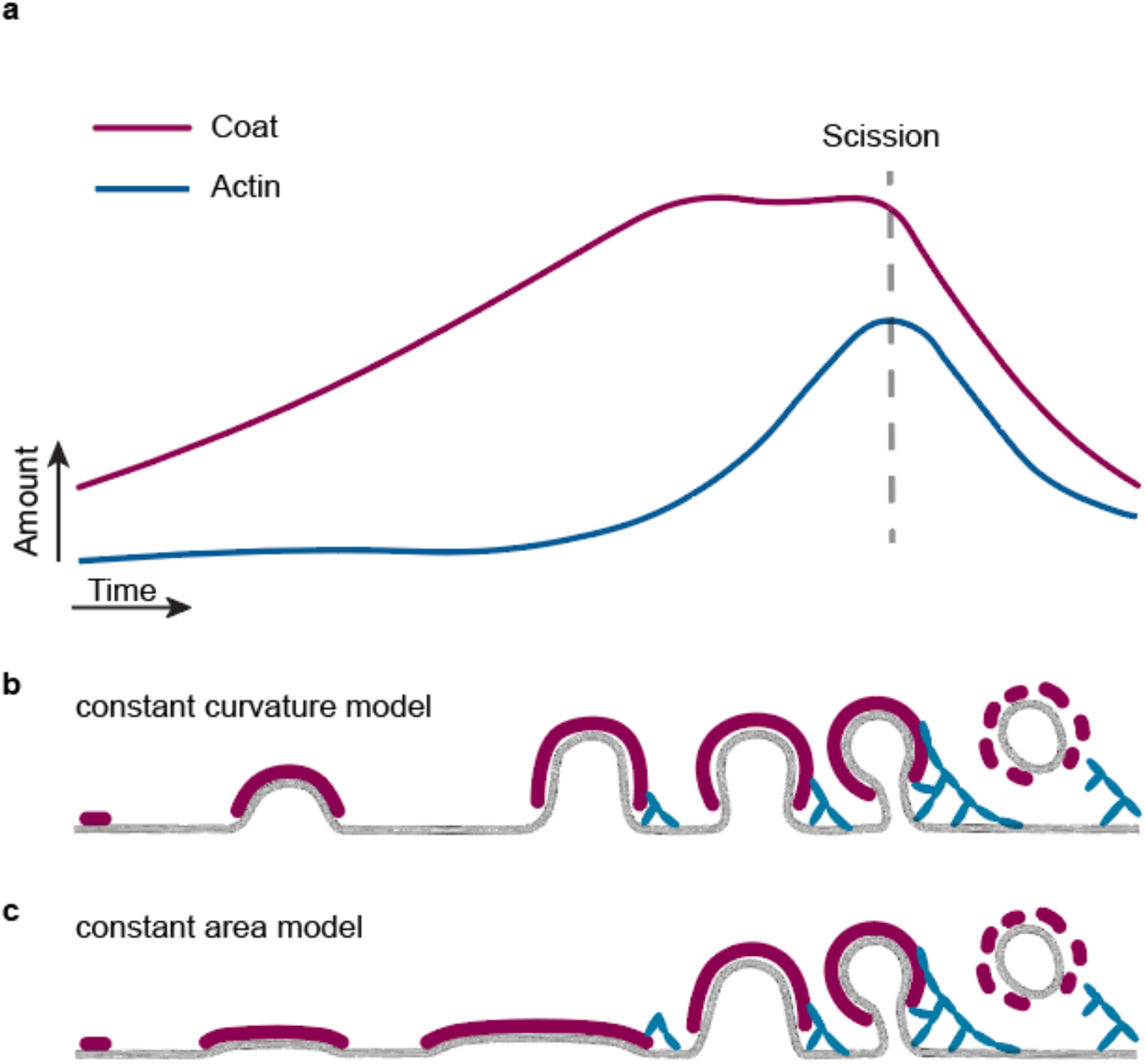
Constant curvature vs constant area models for how clathrin coats assemble at actin-positive CME sites. **a**, A sketch showing amounts of CME coat and actin module proteins at actin-positive CME sites as a function of time based on the data in Fig. 4b. The CME coat is assembled to its maximum area around the time of actin assembly initiation. **b**, Schematic representation of constant curvature model for CME. CME coat assembles during invagination and actin assembles only at deep invaginations. **c**, Schematic representation of constant area model for CME. The CME coat expands to its maximum area first and bends during membrane invagination. In these two different scenarios, actin assembles at CME sites with different curvatures.

